# AnnFlux: object-conditioned neural stochastic differential equations for single-cell perturbation dynamics

**DOI:** 10.64898/2026.09.01.748703

**Authors:** Heesun Choi, Gaeun Byeon, Hayoon Park, Jongseo Park, Sungbin Lim, Joon-Yong An

**Affiliations:** Department of Integrated Biomedical and Life Science, Korea University, Seoul, 02841, Republic of Korea; National Research Laboratory for Convergence Degradation Biology, Korea University, Seoul, 02841, Republic of Korea; Interdisciplinary Major Program in Targeted Degradation-based Innovative Therapeutics, Korea University, Seoul, 02841, Republic of Korea; L-HOPE Program for Community-Based Total Learning Health Systems, Korea University, Seoul, 02841, Republic of Korea; School of Biosystem and Biomedical Science, College of Health Science, Korea University, Seoul, 02841, Republic of Korea; School of Health and Environmental Science, College of Health Science, Korea University, Seoul, 02841, Republic of Korea; Department of Statistics, Korea University, Seoul, 02841, Republic of Korea; Korea Brain Research Institute (KBRI), Daegu 41062, Republic of Korea

## Abstract

Single-cell perturbation profiling measures responses to genetic and chemical interventions, yet most models learn a static map, ignoring how populations move over time and how perturbations combine. AnnFlux, an object-conditioned stochastic differential equation, learns a drift field in latent cell-state space. Conditioning on the perturbing object makes the field queryable one object at a time, yielding per-object drifts comparable across genes and drugs. By learning a drift field tailored to each perturbation context, it interpolates a held-out timepoint in an epithelial-mesenchymal transition time course and predicts unseen perturbations. Beyond point estimates, AnnFlux improves distributional fidelity and predicts responses to held-out perturbation combinations. An IFN-response signature predicted by AnnFlux was associated with TLS proximity in an independent pan-cancer spatial atlas. This framework maps perturbation-driven cell-state evolution as continuous trajectories and represents unseen perturbations using prior-knowledge embeddings.

## Introduction

Predicting how cells respond to perturbations is central to mechanism discovery and therapeutic design.^1,2^ The space defined by genes, drugs, their combinations and biological contexts is far larger than any screen can measure, so models must predict responses that have not been observed directly. Compounding this challenge, several studies have shown that perturbation responses are heterogeneous at single-cell resolution, due to the different basal cellular states.^2–4^ Therefore, a model should learn how an unseen perturbing object moves a population from its initial state distribution through cell-state space, rather than only estimating an average endpoint. Single-cell RNA sequencing (scRNA-seq) further shows that perturbation responses vary across cell states and unfold over time rather than collapse to a fixed endpoint.^2,3,5^

A useful model must therefore generalize to novel perturbations, combinations and contexts, and link observed variation to the perturbing agent, its trajectory and the programs it engages. Most computational models instead predict a single endpoint. Autoencoders (CPA, chemCPA), a graph neural network over gene knowledge graphs (GEARS), learned perturbation-to-expression maps (scGPT and Scouter), conditional generative models that map a perturbation representation to a response distribution (PerturbNet) and optimal-transport models such as Conditional Monge (CMonge) return post-perturbation profiles or distributions and generalize to unseen genes, drugs or combinations,^6–12^ but each treats the response as a static shift rather than a trajectory through cell-state space.

RNA-velocity methods such as velocyto and scVelo learn velocity fields that order cells along a trajectory but are not calibrated to real time.^13,14^ Trajectory generators such as PRESCIENT, scDiffEq, and ARTEMIS do carry a source population forward in real time and interpolate held-out timepoints, yet fit one field that is not resolved by the perturbing object.^15–17^ Their predictions therefore cannot be decomposed per object or recomposed for unseen drug or gene combinations. Flow-matching models such as CellFlow and scDFM condition on the perturbing objects and generalize to unseen perturbations and combinations, but their transport is deterministic.^18,19^

Here we introduce AnnFlux, an object-conditioned neural stochastic differential equation (SDE) that closes this gap. AnnFlux learns an object-conditioned drift field that guides stochastic transport in a latent cell-state space, carrying control cells toward their perturbed state, and unifies endpoint prediction and dynamics in one conditional field. Such predictions can generate counterfactual response signatures for comparison with patient-derived and tissue-resolved molecular atlases.

## Results

### AnnFlux models perturbation response as object-conditioned drift-field dynamics

AnnFlux learns a drift field over cell state, integrating stochastic trajectories from control cells and matching their endpoint distribution to the measured perturbed population (**Fig. 1a, Methods**). In the latent SDE, this field is the drift term, a vector field over the latent space. Within each dataset, AnnFlux conditions the field on the perturbing objects, genes or drugs applied singly or in combination, rather than fitting a separate field for each perturbation. Each object carries a pretrained embedding from a prior-knowledge network (PKN), STRING for genes and LINCS for drugs^20,21^, so that biologically related objects sit near one another, and the field extends to perturbations never seen in training (**Methods**).

**Fig. 1.**
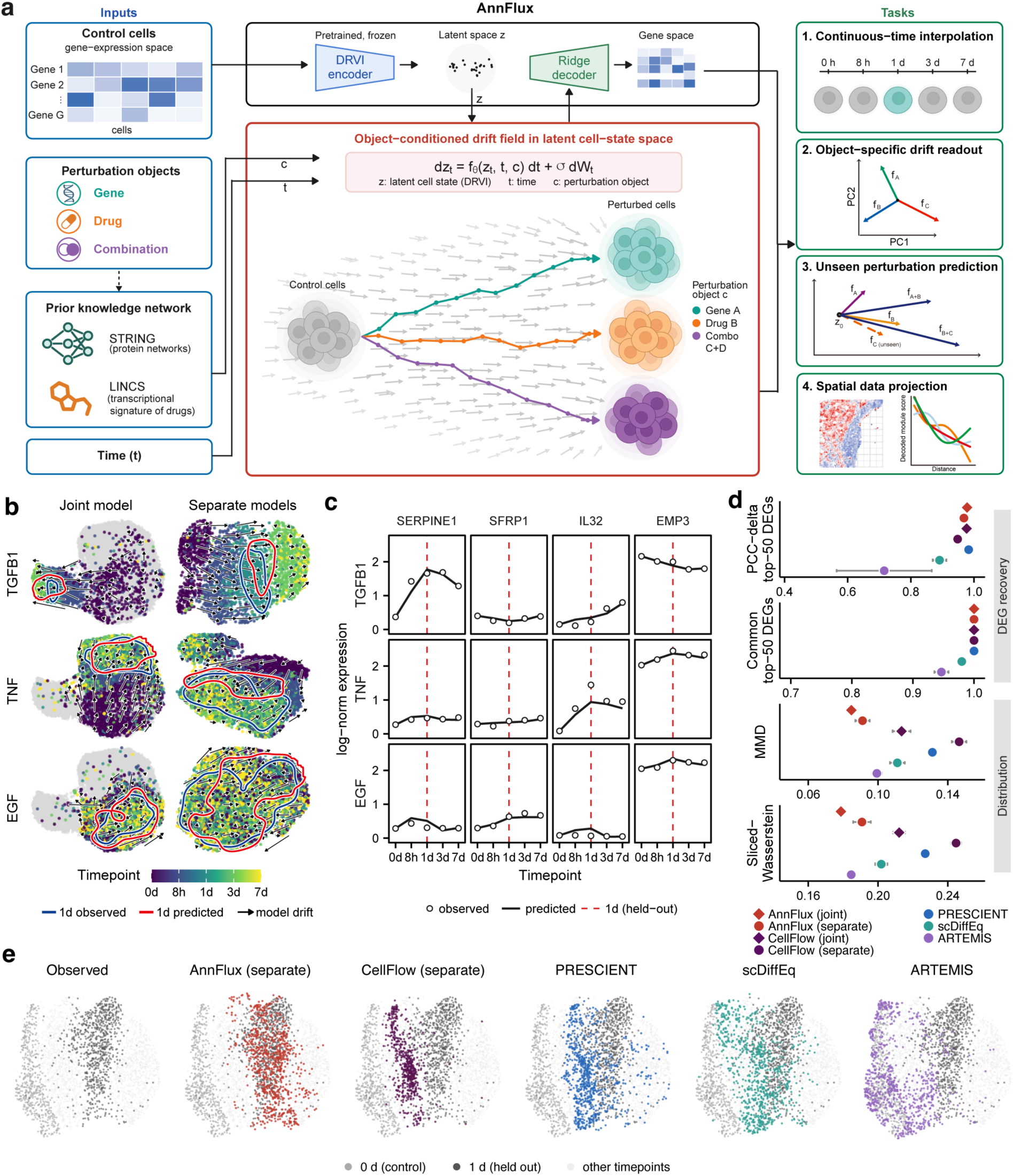
AnnFlux framework and held-out time-course recovery in A549 EMT. **a,** AnnFlux schematic: object-conditioned SDE drift field f_θ_(z, t, c) in a DRVI latent, decoded to gene space. **b,** Held-out 1 d recovery for TGF-β1, TNF and EGF. The joint model is shown in a shared-DRVI UMAP (left), and each separate model in its independently fitted inducer-specific DRVI UMAP (right); observed (blue) and predicted (red) 1 d contours. n = 788, 1,651 and 1,955 observed 1 d cells for TGF-β1, EGF and TNF. **c,** Marker-gene trajectories per inducer, observed versus predicted with 1 d held out; three inducer-defining markers (SERPINE1, SFRP1, IL32) plus EMP3, a pan-inducer EMT gene. **d,** 1 d holdout benchmark on TGF-β1: AnnFlux and CellFlow (each joint and separate; diamonds mark joint models and circles mark per-inducer models) versus PRESCIENT, scDiffEq and ARTEMIS. Higher values indicate better DEG recovery, whereas lower MMD and Sliced-Wasserstein values indicate better distributional fidelity; mean ± s.d. over three training seeds. **e,** Held-out 1 d TGF-β1 recovery in a reference PCA-UMAP fitted to all measured timepoints (n = 577 control cells at 0 h, 788 observed cells at 1 d). Separate panels show the observed population and five models (AnnFlux separate, CellFlow separate, PRESCIENT, scDiffEq and ARTEMIS). In every panel, other measured timepoints are palest gray, the 0 h control is mid-gray and the observed 1 d population is dark gray. Each model’s predicted 1 d population is colored by model. Panels b, c and e show predictions from seed 42; panel d reports mean ± s.d. across seeds 42, 1024 and 2026.

We first asked whether this shared representation could predict an unseen timepoint. In the Cook time course, A549 cells were exposed to three epithelial-mesenchymal transition (EMT) inducers, TGF-β1, TNF and EGF.^3^ Cells were sampled at five timepoints, 0 h, 8 h, 1 d, 3 d and 7 d. Because the AnnFlux drift field is defined as a function of time, it can be evaluated at an unobserved intermediate timepoint. To test this, we withheld the 1 d sample and reconstructed each inducer’s cell-state distribution from 0 h controls. We compared joint and separate modeling strategies. A single joint model represented the three inducer drifts in one shared DRVI latent space. Each separate model instead used an independently fitted DRVI and learned dynamics only from its corresponding inducer (**Fig. 1b**). The joint model also resolved the inducers at the level of individual genes, recovering both the divergent trajectories of each inducer’s defining marker and the shared dynamics of EMP3, a marker active under all three, in agreement with the withheld 1 d measurements (**Fig. 1c**).^3^

Benchmarked against the trajectory and neural SDE models PRESCIENT, scDiffEq, ARTEMIS and the flow-matching model CellFlow under a common protocol, AnnFlux reconstructed the held-out 1 d population faithfully (**Fig. 1d**, **Table 1**). Each prediction was scored for differential-expression recovery using PCC-delta top-50 DEGs and Common top-50 DEGs. Distributional fidelity was assessed using MMD and Sliced-Wasserstein distance. Joint and separate AnnFlux achieved PCC-delta top-50 DEGs of 0.978 and 0.968, respectively, compared with 0.983 for PRESCIENT and 0.977 for CellFlow (joint). Common top-50 DEGs reached 1.000 for AnnFlux, CellFlow and PRESCIENT, compared with 0.980 for scDiffEq and 0.947 for ARTEMIS. The joint AnnFlux model recorded the lowest MMD and Sliced-Wasserstein distance (0.085 and 0.179); independent separate models achieved 0.091 and 0.191. For TGF-β1, the shared joint model had equal or better mean scores than separate training across these four metrics. In uniform manifold approximation and projection (UMAP) space, AnnFlux captured the location and spread of the observed target population (**Fig. 1e**).

**Table 1.** Benchmark of A549 EMT time-course interpolation at the 1-day holdout under TGF-β1.

| Model | PCC-delta top-50<br>DEGs | Common top-50 DEGs | MMD | Sliced-Wasserstein |
| --- | --- | --- | --- | --- |
| AnnFlux (separate) | $0.968 \pm 0.006$ | $1.000 \pm 0.000$ | $0.091 \pm 0.004$ | $0.191 \pm 0.005$ |
| AnnFlux (joint) | $0.978 \pm 0.003$ | $1.000 \pm 0.000$ | $0.085 \pm 0.001$ | $0.179 \pm 0.003$ |
| CellFlow (separate) | $0.948 \pm 0.004$ | $1.000 \pm 0.000$ | $0.146 \pm 0.004$ | $0.245 \pm 0.003$ |
| CellFlow (joint) | $0.977 \pm 0.009$ | $1.000 \pm 0.000$ | $0.113 \pm 0.005$ | $0.212 \pm 0.003$ |
| PRESCIENT | $0.983 \pm 0.005$ | $1.000 \pm 0.000$ | $0.131 \pm 0.001$ | $0.227 \pm 0.001$ |
| scDiffEq | $0.891 \pm 0.022$ | $0.980 \pm 0.000$ | $0.111 \pm 0.004$ | $0.202 \pm 0.003$ |
| ARTEMIS | $0.716 \pm 0.151$ | $0.947 \pm 0.012$ | $0.099 \pm 0.003$ | $0.185 \pm 0.002$ |

### AnnFlux resolves inducer-specific EMT programs by object-wise drift decomposition

AnnFlux conditions the drift on the perturbing object, so the trained field can be queried one inducer at a time to read out its specific drift. We used this to ask whether a single joint A549 model recovers the inducer-specific EMT biology reported for this system.^3^ The model was trained under a cell holdout rather than the timepoint holdout used above (**Methods**). Averaging the per-inducer drift over 0 h cells from the held-out split across three training seeds gave one latent drift vector per inducer. The mean drift magnitude was 37 ± 3 latent units for TGF-β1, 29 ± 4 for TNF and 20 ± 3 for EGF, with the spread giving the seed-to-seed standard deviation (**Fig. 2a**). Decoding these drifts into gene space and projecting them onto the first two principal components gave each inducer its own drift direction (**Fig. 2b**).

**Fig. 2.**
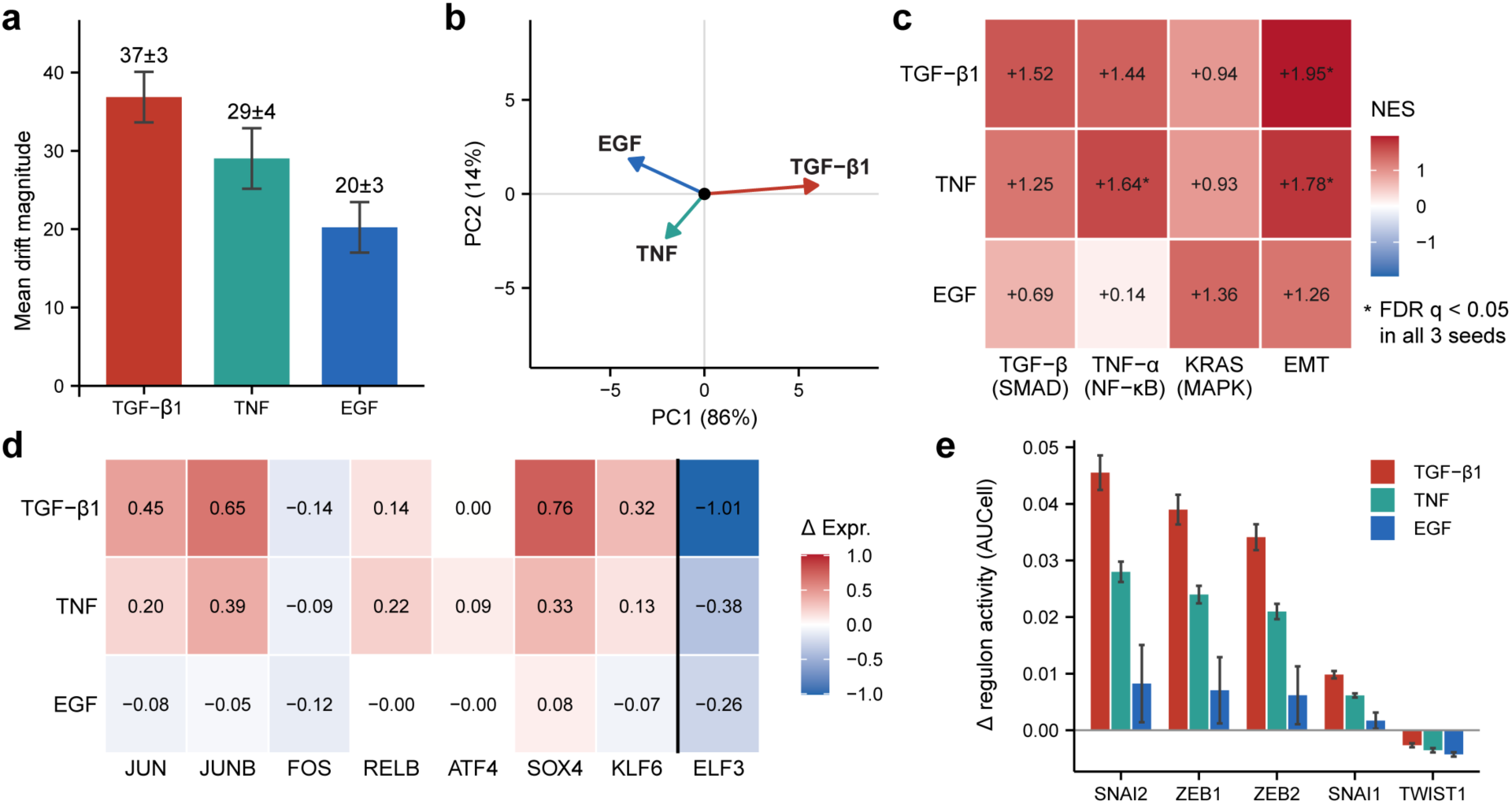
Object-wise drift decomposition resolves inducer-specific EMT programs in A549 cells. **a,** Mean drift magnitude per inducer over 400 held-out control cells at 0 h (mean ± s.d. over three training seeds). **b,** Principal-component projection of the decoded gene-space drift directions for the three inducers (PC1, 86%; PC2, 14%). **c,** Pathway specificity of the decoded drift. Each cell shows the normalized enrichment score (NES) from Hallmark gene-set enrichment analysis (GSEA) for one inducer-pathway pair; matched pairs lie on the diagonal. n = 300 held-out control cells at 0 h. Values are mean NES across three training seeds. Asterisks indicate GSEA FDR q-values below 0.05 in all three seeds (1,000 gene-set permutations per seed). **d,** Decoded change in expression for EMT-associated TFs, grouped as mesenchymal and immediate-early versus the epithelial factor ELF3. **e,** Change in EMT master-regulon activity from 0 h to the drift-driven endpoint, scored by AUCell on the CollecTRI network, for five regulators (SNAI2, ZEB1, ZEB2, SNAI1, TWIST1) per inducer (mean ± s.d. over three seeds).

To test how changing the inducer token reshaped the decoded pathway program, we swapped inducer tokens on a fixed pool of 0 h cells from the held-out split and re-scored each decoded drift against canonical signaling pathways. Three pathways served as diagnostic axes, one per inducer, with Hallmark EMT as the biological readout. On every diagnostic axis, the matched inducer scored highest of the three, separating TGF-β/SMAD, TNF-α/NF-κB and KRAS/MAPK programs by inducer identity (normalized enrichment score, NES = 1.52, 1.64 and 1.36 for TGF-β1, TNF and EGF, respectively). The EMT axis further separated the inducers, with strongest enrichment for TGF-β1 and TNF (NES = 1.95 and 1.78, respectively; FDR q < 0.05 in all three seeds). EGF showed weaker EMT enrichment that was not significant in any of the three seeds (mean NES = 1.26), consistent with the report that EGF does not induce an EMT program in A549 cells (**Fig. 2c**).^3^ At the transcription-factor (TF) level, the decoded expression changes reproduced the A549 pattern reported in the Cook time course: AP-1 (JUN, JUNB), SOX4 and KLF6 rose while the epithelial factor ELF3 fell.^3^ TGF-β1 and TNF shifted these factors in the mesenchymal direction, whereas under EGF the shifts were weak (**Fig. 2d**). The same pattern held for the core EMT master regulators. *SNAI2*, *ZEB1*, *ZEB2* and *SNAI1* rose most under TGF-β1, less under TNF and least under EGF, while TWIST1 did not rise under any inducer (**Fig. 2e**). In the Cook time course, SNAI2 was likewise detected among the canonical EMT factors while TWIST1 was not.^3^ Across these readouts, object-wise decomposition resolved a separate drift for each inducer, and each drift was consistent with the EMT biology reported for this system, so the trained field gives a program-level readout for every perturbing object.

### AnnFlux predicts perturbations across genetic and chemical screens

We next evaluated unseen-perturbation prediction across genetic and chemical screens. Each perturbation was held out, and all models were trained under a common protocol (Methods).

We first benchmarked single-perturbation prediction across three genetic and one chemical perturbation datasets, comparing AnnFlux against baseline models and a training-mean null. Across the three genetic screens AnnFlux gave the lowest distribution distances, outperforming every learned baseline per perturbation (win rates 0.89 to 1.00 for Sliced-Wasserstein and 0.70 to 1.00 for MMD; Methods). Point-estimate baselines, including the training-mean null (trainMean), captured the mean shift but not the cell-to-cell variance. On Norman, AnnFlux therefore reduced MMD relative to trainMean (0.048 versus 0.146). For Common top-50 DEGs, AnnFlux led the learned baselines on Norman and Frangieh, whereas response-direction recovery by PCC-delta top-50 DEGs was screen-dependent and favored scGPT in some settings (**Fig. 3a**; Common top-50 = 0.81 versus 0.78 for GEARS on Norman and 0.82 versus 0.64 for scGPT on Frangieh). The chemical screen showed the converse trade-off, with AnnFlux giving the best differential-expression recovery on sci-Plex3 (Common top-50 DEGs = 0.82, versus 0.77 for CMonge and 0.59 for chemCPA). The optimal-transport baseline CMonge nonetheless led on distribution distances, with AnnFlux second and ahead of chemCPA (**Fig. 3b**).

**Fig. 3.**
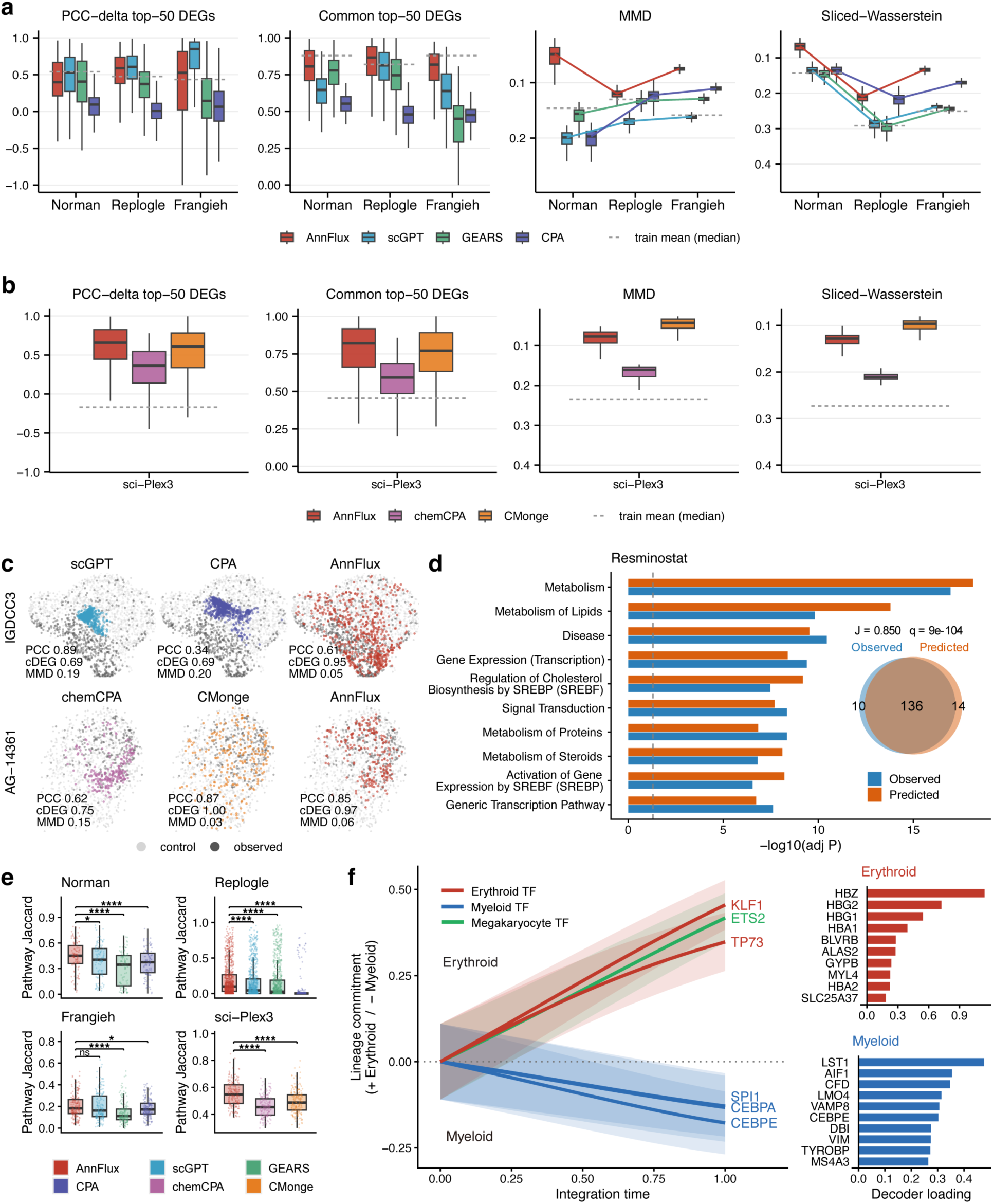
A screen-specific AnnFlux model predicts held-out perturbations in genetic and chemical screens. **a,** Single-gene benchmark (Norman, Replogle, Frangieh). Per-perturbation distributions of four scores. PCC-delta top-50 DEGs and Common top-50 DEGs, differential expression. MMD and Sliced-Wasserstein, distribution overlap with the y axis reversed so higher is better. Lines, per-model medians. Dashed line, training-mean null. Boxes, median and interquartile range (IQR). Whiskers, 1.5× IQR. Models, AnnFlux, scGPT, GEARS, CPA. n = 104, 1,695 and 178 perturbations. The two differential-expression panels are restricted to perturbations with at least ten significant differentially expressed genes (n = 103, 1,289 and 155). AnnFlux versus each baseline on MMD and Sliced-Wasserstein, two-sided Wilcoxon with Benjamini-Hochberg correction (all q < 1×10⁻¹⁷). **b,** As in a, for the single-drug screen (sci-Plex3, n = 188; n = 164 for the differential-expression panels), comparing AnnFlux, chemCPA and CMonge. **c,** Distribution-overlay UMAPs for held-out IGDCC3 (gene, 613 cells) and AG-14361 (drug, 233 cells). Light gray, control. Dark gray, observed. Color, model prediction. Insets, per-perturbation PCC-delta top-50 DEGs, Common top-50 DEGs and MMD. **d,** Pathway recovery for held-out Resminostat. Observed versus predicted enriched Reactome pathways (top 10, -log₁₀ adjusted P, dashed line at FDR = 0.05). Area-proportional Venn with Jaccard index and hypergeometric q. **e,** Pathway-set Jaccard across held-out perturbations with at least five significantly enriched observed pathways, per screen (n = 104/894/176/188 for Norman/Replogle/Frangieh/sci-Plex3). AnnFlux versus each baseline, one-sided paired Wilcoxon (asterisks mark q < 0.05, 0.01, 0.001 and 1×10⁻⁴, ns not significant). **f,** Held-out lineage commitment along AnnFlux integration time t for six Norman TFs (band, 95% CI). Erythroid (KLF1, TP73) and megakaryocyte (ETS2) drivers rise, myeloid drivers (SPI1, CEBPA, CEBPE) fall. Right, top decoder-loading genes per program.

In single-cell UMAP overlays of a held-out gene knockout and a drug, the point-estimate baselines (scGPT, CPA, chemCPA) produced substantially contracted predicted distributions. CMonge, which matches distributions directly, spread its predictions but was more diffuse than the observed population. AnnFlux instead spread cells across the observed distribution (**Fig. 3c**).

The predictions also reproduced downstream biological programs. For the held-out histone deacetylase (HDAC) inhibitor Resminostat, the pathways in the Reactome 2022 pathway collection that were enriched in the predicted response matched the observed profile, sharing 136 pathways at a Jaccard of 0.85 (**Fig. 3d**). Across held-out perturbations, AnnFlux showed the highest overlap in each of the four screens, although pathway recovery was limited for all models on Replogle (**Fig. 3e**). Because AnnFlux predicts a trajectory rather than a static endpoint, its predictions also show how a perturbation commits cells along that trajectory. We integrated the drift field from control cells for six held-out Norman TFs. Erythroid-megakaryocytic drivers (*KLF1*, *TP73*, *ETS2*) and myeloid drivers (*SPI1*, *CEBPA*, *CEBPE*)^22,23^ drove cells toward opposite ends of an erythroid-myeloid axis (**Fig. 3f**).

This object-generalization setting makes the prior-knowledge network most relevant in large Perturb-seq screens such as Norman, where held-out gene perturbations are sparse but biologically coupled through shared regulatory and signaling networks. Ablating the PKN produced modest overall effects. The paired effect (Cohen’s d^z^, oriented so that positive values indicate better performance with PKN, paired on held-out perturbations) was positive in 43 of 52 dataset × regime × metric cells and reached q < 0.05 in 23. Effects were modest and consistent rather than large: the median |d^z^| was 0.23, and the effect on the largest screen was 0.43 on Replogle PCC-delta top-50 DEGs (q < 0.001). The prior also organized learned drift into known gene modules, for which the recovery area under the precision-recall curve (AUPRC) rose from 0.15 to 0.28 for the HUGO Gene Nomenclature Committee (HGNC) gene family and from 0.42 to 0.78 for TF lineage. Together, AnnFlux captured perturbed-population distributions while maintaining competitive differential-expression recovery across genetic and chemical screens.

### AnnFlux predicts combinatorial perturbations and their interaction structure

We next asked whether AnnFlux extends from single to combinatorial perturbations, where the central difficulty is generalization to combinations that were never observed (**Fig. 4a**). We defined three regimes of increasing difficulty according to how many of a combination’s two constituent single perturbations were seen during training: both (“2 seen”), one (“1 seen”), or neither (“0 seen”) (**Methods**).

**Fig. 4.**
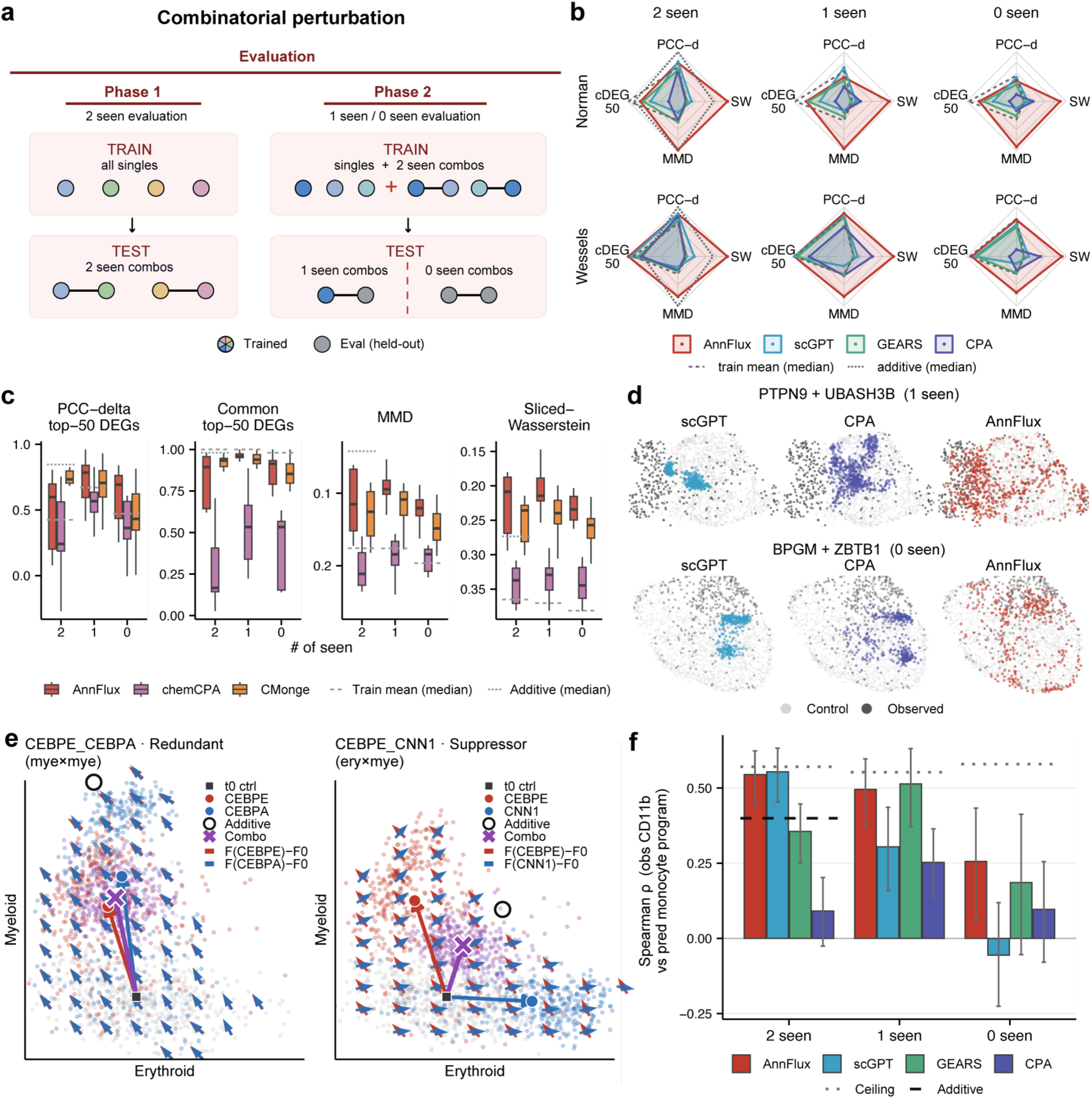
AnnFlux predicts combinatorial perturbations and recovers their interaction structure. **a,** Schematic of the held-out combinatorial evaluation across the 2, 1 and 0 seen regimes. **b,** Radar plots of prediction accuracy on the genetic combinatorial datasets (Norman and Wessels rows, 2, 1 and 0 seen columns). Axes, PCC-delta top-50 DEGs, Common top-50 DEGs, MMD and Sliced-Wasserstein. Axes are rescaled within each dataset, so radial position denotes relative standing and a larger envelope is better. Underlying medians are computed per held-out combination after averaging over the available evaluation seeds. Dashed and dotted contours, training-mean and additive baselines. Models, AnnFlux, scGPT, GEARS, CPA. n = 131/107/46 (Norman) and 158/129/63 (Wessels). **c,** Box plots of the same metrics on drug combinations. Boxes, median and IQR. Whiskers, 1.5× IQR. Dashed and dotted lines, training-mean and additive medians. Models, AnnFlux, chemCPA, CMonge. n = 7/19/11. **d,** UMAP scatter of predicted versus observed single-cell distributions for two held-out Norman combinations. Light gray, control. Dark gray, observed. Color, model prediction (scGPT, CPA, AnnFlux). **e,** Latent drift fields for redundant CEBPE + CEBPA and suppressor CEBPE + CNN1 combinations along erythroid and myeloid programs. Small arrows show baseline-subtracted single-perturbation drifts, F(A) - F0 and F(B) - F0; the open circle and cross mark additive and predicted combination endpoints. Two representative pairs are shown for seed 42, fold 0. **f,** Spearman correlation between observed CD11b and the predicted monocyte program. Bars, mean over three seeds. Error bars, bootstrap 95% confidence interval over combinations (10,000 resamples, drawn once per stratum and shared across series so the series are paired), not the seed standard deviation. Dotted line, ceiling. Dashed line, additive baseline. Models, AnnFlux, scGPT, GEARS, CPA. n = 142 per seed/129/63. Exposure strata in b, c and f are not disjoint; they regroup the same held-out combinations (131 Norman, 158 Wessels, 25 drug).

Across all three regimes, AnnFlux gave the lowest distribution distances of the learned models. AnnFlux achieved high per-combination win rates against every learned baseline (Sliced-Wasserstein and MMD win rates 0.98 to 1.00 for genetic screens and 0.82 to 1.00 for drug combinations). On Norman at 2 seen, this corresponded to low absolute distribution distances (median MMD = 0.038, median Sliced-Wasserstein = 0.060). On differential-expression recovery it matched the strongest point-estimate baselines, scGPT, GEARS and CPA, with Common top-50 DEGs of 0.84 on Norman and 0.98 on Wessels. From 2 seen to 0 seen, AnnFlux held the identity of the responding genes and its distributional advantage. On Norman, Common top-50 DEGs stayed at 0.84 and its win rate against every learned baseline remained 1.00 on both distance metrics. PCC-delta top-50 DEGs fell for all four models on Norman. The decline was smaller on Wessels, where AnnFlux fell from 0.88 to 0.83 and stayed ahead of scGPT, GEARS and CPA at every regime (**Fig. 4b**). The same pattern held for drug combinations. AnnFlux again gave the lowest distribution distances across regimes. On differential-expression recovery it was on par with CMonge, which led at 2 seen, while AnnFlux led at 1 seen and 0 seen, although the small stratum sizes should be considered when interpreting these differences (n = 7-19) (**Fig. 4c**).

These distances have a direct visual counterpart. For the held-out Norman combinations shown, AnnFlux recovered both the mean shift and cell-to-cell variation, whereas scGPT and CPA predictions were more concentrated around their means (**Fig. 4d**). Beyond recovering the population distribution, the object-wise drifts also expose how the component perturbations combine. Because combinatorial perturbations need not be additive, we projected the predicted cell cloud from the Norman study onto curated erythroid and myeloid program axes to examine how the drift field was organized across lineage programs.^22^ For the myeloid TFs *CEBPE* and *CEBPA*^23^, the two drifts align and the pair is redundant. By contrast, the predicted drifts for *CEBPE* and *CNN1*^22^ pointed toward opposing lineage programs. The combination occupied an intermediate position, consistent with its suppressor annotation^8^ (**Fig. 4e**). As an orthogonal cross-modality test in the Wessels Cas13 RNA Perturb-seq screen of chromatin regulators driving monocytic differentiation in THP1 cells^24^, we tested whether the monocyte differentiation program predicted by AnnFlux for held-out gene pairs ranked the combinations in agreement with the paired CD11b readout, an immunophenotypic marker of monocyte differentiation (mean Spearman ρ = 0.545, 0.496, and 0.256 for 2 seen, 1 seen, and 0 seen, respectively; **Fig. 4f**).

### An AnnFlux-predicted IFN-response signature is associated with TLS proximity in a pan-cancer spatial atlas

Having learned drift fields from in vitro perturbation screens and simulations of held-out perturbations, we next asked whether a signature that AnnFlux predicts for held-out perturbations reappears in an unseen data modality. We read a gene-level signature out of the learned drift field for perturbations withheld from training. Recapitulating this predicted signature in independent tissue would indicate that the prediction carries transferable response structure.

Cho et al. reported that immune pathways, including IFN-α and IFN-γ responses, major histocompatibility complex (MHC) class II antigen presentation and inflammatory signaling, were highest near intratumoral TLSs and decreased with distance (high-to-low).^25^ We used an IFN-response signature predicted by AnnFlux in the IFN-γ-stimulated Frangieh melanoma CRISPR-knockout screen.^26^ We read it from the drift predicted for *STAT1* and *JAK2*, two IFN-γ/JAK-STAT nodes withheld from training (**Methods**).^27,28^ We asked whether the drift field predicted for these held-out IFN-γ/JAK-STAT nodes encoded an IFN signaling program and whether the resulting signature reappeared as a spatially organized program across tumor tissues when projected onto an independent pan-cancer spatial atlas of tertiary lymphoid structures (TLSs).^25^

We scored the AnnFlux signature in all 35 sections of the atlas, spanning eight cancers. It was TLS-proximal in 22 of 35 sections and TLS-distal in the other 13, and the curated Hallmark IFN-γ signature was TLS-proximal in 20. The two signatures agreed on 27 sections, and their section-level gradient strengths were correlated (Pearson r = 0.65).

In one kidney section and one liver section, the drift-derived IFN signature was highest next to TLSs and declined with distance (per-spot Pearson r -0.56 and -0.51; **Fig. 5a**). In the same two cancers it ranked among the immune and antigen-presentation programs that met the high-to-low criterion (**Fig. 5b**). Pooled within each cancer, under a stricter three-way classification, Hallmark IFN-γ met the high-to-low criterion in six of the eight cancers, and the AnnFlux signature recovered the same gradient in five of those six (**Fig. 5c**). Colorectal cancer was the exception, declining with distance but not meeting the criterion in a reference-signature-selected subset used as a positive-control comparison. Overall, the predicted signature was associated with TLS proximity in independent tissue, with variation across sections and cancers.

**Fig. 5.**
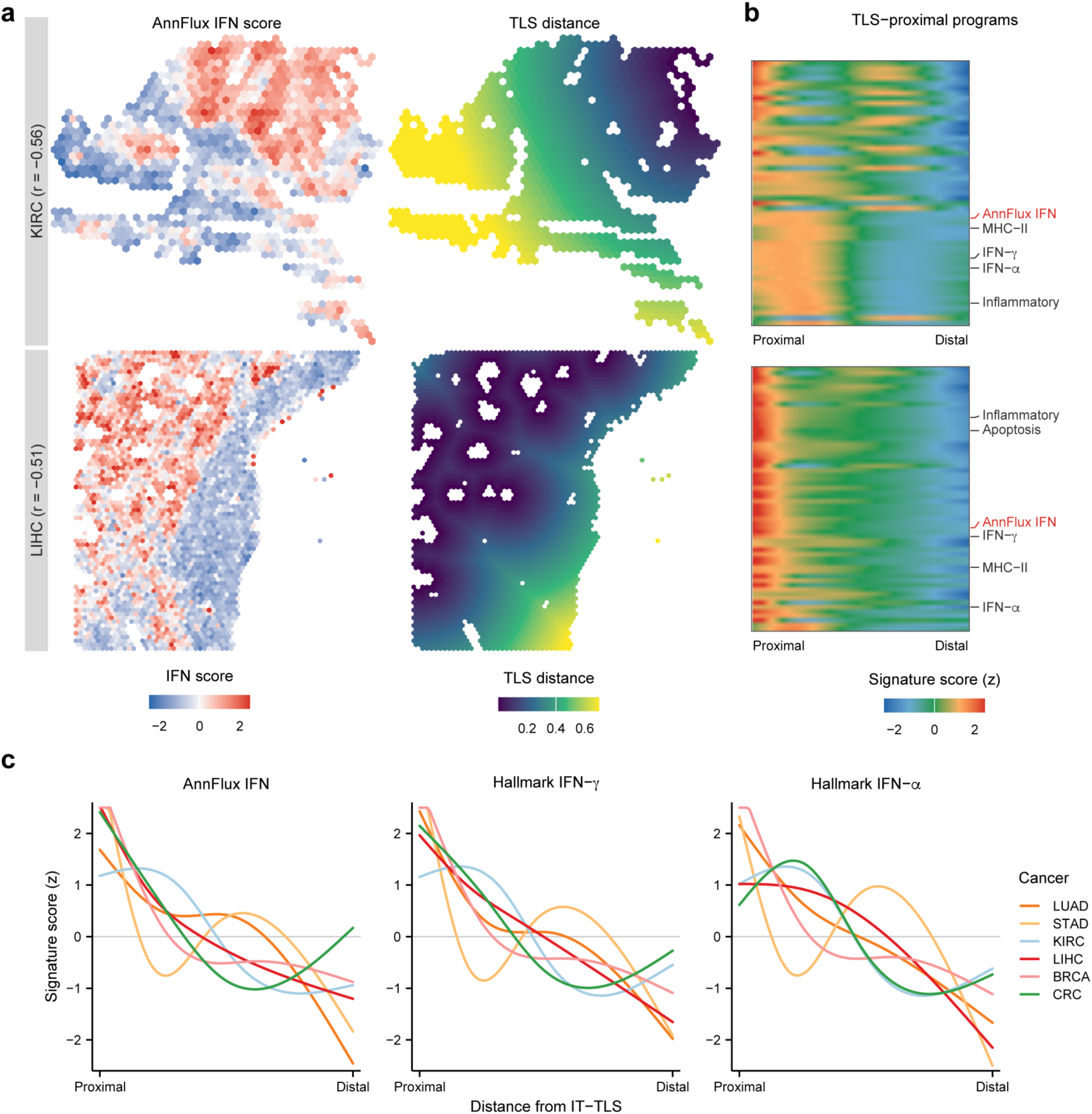
Association between an AnnFlux-derived IFN-response signature and TLS proximity in the Cho spatial atlas. **a,** AnnFlux IFN-response signature (AnnFlux IFN), predicted under perturbation-holdout, projected onto two Cho atlas Visium sections. One clear-cell renal carcinoma (KIRC, ccRCC S7, n = 1,076 spots) and one liver (LIHC, Liver S9, n = 3,142 spots). Left, AnnFlux IFN score. Right, normalized distance from intratumoral TLSs (IT-TLSs). Per-spot Pearson r, -0.56 (KIRC) and -0.51 (LIHC). **b,** Distance-ordered heatmaps of TLS-proximal programs for the two cancers. Rows, Cho atlas programs with a significant high-to-low gradient, plus AnnFlux IFN. Selected immune and antigen-presentation programs, annotated at right. AnnFlux IFN, red. Color, z-scored signature activity from proximal to distal. **c,** Smoothed distance profiles (natural-spline fits) for AnnFlux IFN, Hallmark IFN-γ and Hallmark IFN-α across the six cancers selected by the Hallmark IFN-γ high-to-low criterion, providing a positive-control comparison independent of AnnFlux. Lines, cancer types. Each fit is evaluated on a 100-point distance grid.

## Discussion

Earlier dynamical models fit a single velocity field that is not resolved by the perturbing object, so their predictions cannot be separated per object or recomposed for unseen inputs.^15–17^ AnnFlux instead conditions the drift on each object, using prior-knowledge embeddings to represent unseen inputs and per-object drifts to aid interpretation.^6,8^

Across both time-interpolation and object-prediction tasks, AnnFlux showed its most consistent advantage on distributional metrics. Its objective explicitly supervises shift magnitude, direction and population shape. In the single-cell UMAP overlays, several baseline predictions were contracted or more diffuse, whereas AnnFlux matched the observed spread. Beyond statistical benchmarks, downstream biological validations of the predicted drift fields recovered known response programs, indicating that object-conditioned dynamics can add biological interpretability to latent representations such as DRVI by linking perturbing agents to downstream programs. For combinations, AnnFlux conditions the same learned drift field on paired perturbing objects, predicting non-additive population responses and exposing interaction structure for unseen pairs. Trained on RNA from single perturbations alone, AnnFlux ordered held-out gene pairs by predicted differentiation, and that order matched the CD11b protein measured on the same cells. Extending these drift-derived outputs to human tumor tissue, a signature predicted for held-out interferon-pathway genes tracked distance from TLSs across an independent pan-cancer spatial atlas.^25^ This link between controlled perturbation screens and tissue-resolved disease programs provides a basis for prioritizing context-specific response programs for downstream experimental and translational studies.

However, several limitations remain. Differential-expression direction was dataset- and response-dependent. On Norman and Frangieh, the training-mean null exceeded AnnFlux on at least one top-50 DEG metric. AnnFlux gave the lowest distribution distances on the three genetic screens, whereas CMonge achieved lower distances on sci-Plex3. Direction recovery weakened as source and target populations became more similar. Several model-configuration comparisons reused the benchmark holdout partitions and are therefore interpreted as retrospective sensitivity analyses rather than independent validation. Generalization to new objects depends on prior coverage, and sparsely covered genes or drugs use less informative out-of-vocabulary embeddings. AnnFlux is currently trained separately for each dataset and does not jointly model multiple doses or cell lines. Extending the conditioning scheme across doses, cellular contexts and datasets will be important for broader use. The spatial findings are observational and may partly reflect local cell-type composition, which future analyses should account for.

By assigning an interpretable drift to each perturbing object, AnnFlux links endpoint prediction to testable hypotheses about response direction, magnitude and interaction geometry. This object-conditioned formulation could support virtual-cell models that simulate responses to unmeasured interventions in silico.^29^

## Methods

### Training, pairing and inference

AnnFlux is trained by simulating its dynamics from source to target and matching the simulated endpoint population to the observed one. Because single-cell measurement is destructive, the true source-to-target correspondence is unobservable. The pairing used in training is therefore a surrogate. Within each training batch, target cells were re-paired to source cells by an entropic optimal-transport coupling computed separately within each group of identical conditioning, that is, the same objects, cell line and time interval, on a squared-Euclidean cost (Sinkhorn, ε_OT_ = 0.05, 50 iterations).^30^ The assignment was drawn from the coupling rather than taken at its maximum, so the pairing is resampled each epoch. The object-conditioned latent stochastic differential equation,

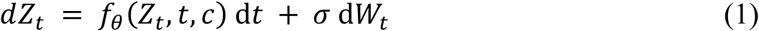

with drift *f_θ_*(*z*, *t*, *c*) and Brownian motion *W_t_* of a constant-noise scale σ = 0.5, is advanced from a source cell by Euler-Maruyama integration (equation (2)), where *t*_src_ and *t*_tgt_ are the source and target times of the pair being integrated and K is the number of integration steps. The state *Z*_t_ takes values in R^32^, the cell latent space. The drift maps a state, a normalized time *t* ∈ [0, 1] and the object-conditioning vector *c* to a vector in R^32^. The scalar time is represented using a 64-dimensional sinusoidal embedding, and the drift is parameterized by an AdaLN-Zero residual network. Training uses K = 20. At inference, the number of integration steps is scaled with the normalized interval length, using approximately 30 steps per unit time. Specifically, K = max(2, ⎣30(*t*_tgt_ − *t*_src_)⎦) for the time course, where *t*_tgt_ − *t*_src_ denotes the normalized interval length. A minimum of two steps is used for short intervals. For the screens, the full normalized interval from 0 to 1 is integrated using K = 30.

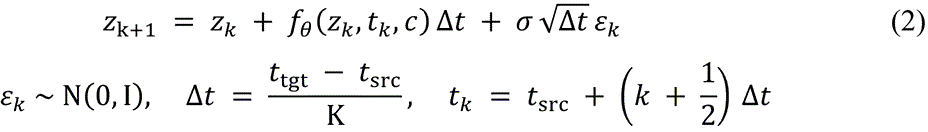

The simulated endpoint population is matched to the optimal-transport-paired target by a grouped, weighted objective (equations (3-7)),

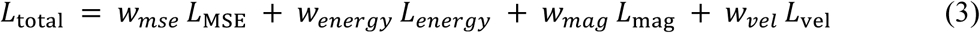

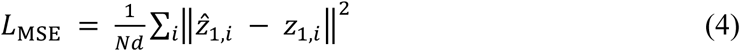

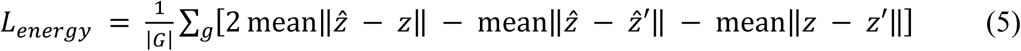

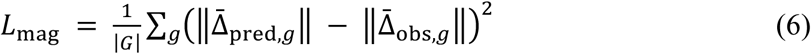

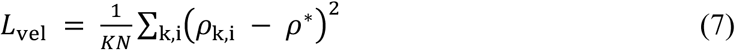

where ẑ₁ is the SDE endpoint integrated from a source cell z₀ and z₁ its optimal-transport-matched target; z′ and ẑ′ are independent samples within a perturbation group g; the group mean shift from control is the mean of a group minus the mean of its control cells; |G| is the number of perturbation groups, N the number of paired cells, K the number of integration steps and σ = 0.5 the fixed noise scale; c is the object-conditioning vector obtained by pooling the perturbation’s object tokens (see Prior-knowledge object encoder). Fallback term weights for unregistered datasets are (w_mse_, w_energy_, w_mag_, w_vel_) = (0.5, 1.0, 1.0, 0.1); in the selected runs the energy and velocity weights are held at 1.0 and 0.1 while the mean-squared-error and magnitude weights are set per dataset and task. The distribution term L_energy_ is the energy distance.^31^ The velocity-ratio term L_vel_ regularizes the per-step drift-to-noise ratio ρ = ǁfθ(zk, tk, c)ǁ Δt / (σ √Δt √d) toward a target ρ* = 2.5, where d = 32 is the latent dimension and the ratio is averaged over integration steps and cells. Because σ is fixed rather than learned, this term sets the scale of the drift relative to the diffusion.

The energy term was computed within grouped batches of 256 cells. The sampler packs as many perturbations per batch as are available up to sixteen and splits the batch evenly among them, so the perturbation screens see sixteen perturbations of sixteen cells while the three-inducer time course sees three groups of about eighty-five. Training used AdamW with task-specific learning rates, a weight decay of 1 × 10⁻⁴ and gradient clipping at a norm of 1.0.^32^ Except for the Cook joint models, the learning rate followed a 10-epoch linear warmup and then a cosine decay. Early stopping generally monitored validation loss with a patience of 30 epochs. For the Cook time-course benchmark, the joint models were trained for 300 epochs, whereas the inducer-specific separate models were trained for 400 epochs. Each Cook run completed its fixed epoch budget and retained the checkpoint with the lowest validation loss. Both Cook regimes used energy, mean-squared-error, velocity-ratio and magnitude weights of 1.0, 0.3, 0.1 and 0, respectively. The ComboSciPlex 2 seen (Phase 1) runs were too small for a validation split, so they instead trained for a fixed 100 epochs and kept the final-epoch weights. At inference, 10 samples were drawn per source cell.

### Noise, integrator and loss settings

AnnFlux was trained with σ = 0.5 and K = 20. We evaluated K ∈ {20, 40, 80, 160, 320} on the Norman single-perturbation, Norman combinatorial and sci-Plex3 screens. No run diverged or produced non-finite latent coordinates, and no metric improved (165 runs).

The energy and velocity-ratio weights, when enabled, were fixed at 1.0 and 0.1, respectively. The target ρ* was fixed at 2.5 for every dataset and task.

### Prior-knowledge object encoder

Each perturbation is represented as a fixed sequence of typed slots, perturbed genes for the genetic screens and compounds for the drug screens. A single perturbation occupies one slot and a combination occupies two, one for each member. Rather than learning object embeddings from the response data alone, the default encoder conditions each object on an external prior-knowledge source: a STRING v12 protein-protein interaction network for genes (Node2Vec embeddings) and LINCS-L1000 transcriptional signatures for drugs.^20,21^ The prior for a source *s* is a fixed feature table 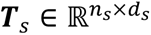 whose rows are precomputed object features, with a map from each object to its row (row 0 denoting out-of-vocabulary). For an object *o* with row *r* in source *s*, the encoder emits a token (equation (8))

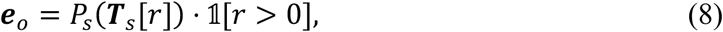

where *P_s_* is a small learned per-source projection into a shared *d* = 64-dimensional embedding space: for genes a LayerNorm followed by a single linear map, and for drugs a LayerNorm followed by a multilayer perceptron with a SiLU nonlinearity. The prior tables ***T****_s_* are frozen throughout training, so the learned parameters capture only how to read the prior rather than the prior itself.

Gene priors are Node2Vec embeddings (dimension 128) of the STRING v12 human network restricted to high-confidence edges (combined score ≥ 700), trained by skip-gram with negative sampling over uniform random walks (walk length 80, 10 walks per node, context window 5, *p* = *q* = 1).^20,33^ Drug priors are LINCS-L1000 consensus signatures.^21^ Object-to-row maps use case-insensitive gene-symbol matching and salt- and synonym-normalized drug-name matching.

Coverage of the priors is incomplete, so out-of-vocabulary objects are resolved in two tiers. At asset-build time a gene absent from STRING is imputed from its nearest STRING neighbors in ESM2 protein-sequence space (top-5 cosine neighbors, minimum cosine 0.3) and a drug absent from LINCS from its nearest neighbors in Morgan-fingerprint Tanimoto space, with gene imputation skipped when STRING coverage of the panel falls below 30% so that too few reliable anchors cannot bias the table.^20,34,35^ An object without a usable prior falls back to a per-object residual. The residual is zero-initialized for a fully out-of-vocabulary object and is learned only if that object appears during training.

For a perturbation with object tokens {***e****_o_*}, a permutation-invariant pooler makes the prediction independent of member order and can represent object-object interactions. Because the objective acts on populations rather than object pairs, performance relative to an additive reference provides an indirect test of non-additive predictive capacity (**Fig. 4f**). The tokens, tagged with a learned slot-type embedding and prepended with a learned pooling query, are passed through a two-layer pre-norm Transformer encoder (*d*_model_ = 128, 4 heads, GELU), and the pooled output is projected back to the object-token space to give a single conditioning vector ***c***. Concatenated with a sinusoidal embedding of the diffusion time *t*, this vector conditions a three-block AdaLN-Zero residual drift field (hidden width 256).^36^ The per-block modulation and the output layer are both zero-initialized, so each block starts as the identity and the initial drift is zero. The influence of the conditioning is learned.

### Datasets and preprocessing

Seven public datasets span the three settings. The time course is Cook^3^ (A549 epithelial-mesenchymal transition under TGF-β1, EGF and TNF). The single-perturbation screens are three genetic (Replogle^5^ CRISPRi and Norman^22^ CRISPRa in K562; Frangieh^26^ CRISPR knockout in IFN-γ-stimulated melanoma) and one chemical (sci-Plex3^2^ in A549). The combination screens are the genetic Norman^22^ and Wessels^24^ and the chemical ComboSciPlex^6^.

Each dataset was normalized to 10⁴ counts and log1p-transformed, then reduced to ∼2,000 highly variable genes^37^ (with raw unique molecular identifier (UMI) counts kept as a separate layer), and encoded by a DRVI variational autoencoder (VAE)^38^ giving a 32-dimensional cell latent. The Cook time course was additionally quality-controlled before normalization, removing cells with fewer than 200 detected genes and genes detected in fewer than 10 cells. DRVI was fitted on raw counts using the default pNB likelihood (a log-space-parameterized negative binomial). For the Cook comparison, the joint model used one DRVI fitted to pooled cells from all three inducers. Each separate model used an independently fitted DRVI containing only its corresponding inducer. We refer to this representation as DRVI NB. The encoder and decoder each had a width of 128. Training used a batch size of 512, a cap of 400 epochs and early stopping. The dynamics were then trained on this fixed latent. AnnFlux uses this DRVI-32 latent representation; each baseline uses the representation defined by its reference implementation.

### Latent representation and decoder

We compared three representations: DRVI NB (raw counts, 32 dimensions), DRVI Gaussian (log-normalized counts, 32 dimensions), and a 50-component principal-component analysis (PCA) of the highly variable gene matrix. For each held-out perturbation we computed a within-representation signal-to-noise ratio (SNR), defined as the L2 norm of the mean shift from control to the perturbed population divided by the square root of the trace of the perturbed-population covariance in that representation. We also summarized how efficiently each representation used its axes by the participation ratio (Σλ)²/Σλ² of the representation-covariance eigenvalues λ, divided by the representation dimensionality. DRVI NB exceeded DRVI Gaussian in SNR in every dataset (two-sided paired Wilcoxon signed-rank test with Benjamini-Hochberg control; q < 0.05 throughout, and q < 0.01 in six). It also used its axes more efficiently than principal-component analysis (q ≈ 0.03, n = 7 datasets), so AnnFlux operates in this latent.

All reported AnnFlux predictions were decoded from the NB DRVI latent space into log-normalized gene space using a closed-form ridge linear readout (λ = 10).

### Tasks, splits and the perturbation-strength gate

AnnFlux and all baselines were evaluated on three tasks using a common metric suite and three training seeds. Five-fold cross-validation was used for the single-perturbation task and combination Phase 2, whereas combination Phase 1 used a 2-seen holdout protocol with seed-specific validation and evaluation partitions. Models were trained separately for each dataset, so a reported model covers the perturbations of its own dataset and not those of the others. The time-course task withholds an interior timepoint of the Cook trajectory and predicts it by transporting 0 h cells to its continuous normalized time t, defined as elapsed hours divided by 168 h so that 7 d mapped to t = 1; the withheld timepoint is 1 d. Two training regimes were compared on this task: a single joint model trained on all three inducers simultaneously, and separate models trained on one inducer each. The joint model operated in a 32-dimensional DRVI latent fitted to all three inducers and used a ridge decoder fitted on the pooled A549 data. Each separate model operated in an independently fitted 32-dimensional DRVI latent and used a ridge decoder based only on its corresponding inducer. The two regimes were compared after decoding their predictions into the same gene-expression space. The single-perturbation task held out individual genes or drugs and predicts the held-out response. The combinatorial task holds out a perturbation pair under two regimes: a combination holdout (Phase 1, 2 seen), in which both constituent singles remain in training, and a member holdout (Phase 2), in which a member and its single are removed so that one (1 seen) or both (0 seen; zero-shot) members are unseen.

Many cataloged perturbations produce no measurable transcriptional change, and scoring them would reward predicting no effect, so perturbations were gated by strength. For each perturbation a sample-size-independent score, SNRd, was computed as the mean of the twenty largest absolute Cohen’s d values relative to control across the highly variable genes. A per-perturbation null floor was estimated from control cells only. Each draw took a pseudo-perturbation group matched to the perturbation’s cell count and a disjoint pseudo-reference group of up to 1,500 cells. SNRd was recomputed over a range of sample sizes (40 repeats) and the 95th percentile interpolated to each perturbation’s cell count. A perturbation with at least 25 cells was classified as detectable when its SNRd exceeded this floor; all others were classified as weak. In the single-perturbation task weak perturbations were kept in training but not evaluated, so reported metrics reflect perturbations with a detectable effect; in the combinatorial task no strength gate was applied, because a weak combination can reflect genuine antagonism, and combinations were included on reliability alone. For sci-Plex3, the input was restricted to the highest-SNRd dose for each drug before gating.

All splits were deterministic. For the single-perturbation and combinatorial tasks, control cells were partitioned once into disjoint training, validation and evaluation pools in proportions of 65/15/20, with the evaluation pool serving as the metric baseline. Randomized split construction used seed 0 for the single-perturbation task and combination Phase 2; for combination Phase 1, the control-cell and combination partitions were redrawn for each training seed (42, 1024 and 2026). Five-fold cross-validation was used for the single-perturbation task and combination Phase 2, with a validation fraction of 0.15. Held-out perturbations were distributed across the five folds by a continuous-effect round-robin on SNRd so that each was evaluated exactly once. Phase 1 instead used a fixed combination holdout with no fold split: each evaluated combination was withheld while both constituent singles remained in training. The numbers satisfying this strict 2-seen criterion were 131/131 for Norman, 158/158 for Wessels and 7/25 for ComboSciPlex. In Phase 2, each held-out gene or drug and its corresponding single were removed from training, and results were pooled by constituent training exposure. The time-course task used two holdout designs. The timepoint holdout excluded the 1 d cells from AnnFlux dynamics training and ridge-readout fitting. These cells were retained for highly variable gene selection and unsupervised DRVI fitting and served as the observed benchmark target. The cell holdout split cells within each inducer-timepoint group into training, validation and test pools in proportions of 70/10/20 using seed 42, with the same source pool and cell cap for every model. This design supplied the joint model used for object-wise drift decomposition, ensuring that the reported drifts were evaluated on cells excluded from training.

### Baselines

AnnFlux was compared against task-appropriate baselines, each run through its own reference implementation under a common evaluation protocol: the same splits, source-cell pool, training seeds and scoring rules within each task. For the time-course task, AnnFlux was compared against four generators that transport a source population forward: PRESCIENT (a potential-driven drift trained with entropic optimal transport in a 50-dimensional PCA space), scDiffEq (a drift-diffusion neural SDE trained with a Sinkhorn divergence in PCA-50), ARTEMIS (a VAE with an unbalanced Schrödinger bridge) and CellFlow (a conditional flow-matching model with entropic optimal-transport couplings in PCA-50).^15–18^ For the perturbation tasks, five learned predictors were trained per fold where applicable: GEARS (a gene-gene Gene Ontology graph network), scGPT (the public whole-human checkpoint fine-tuned per task), CPA (a compositional perturbation autoencoder) and its chemical extension chemCPA (with RDKit molecular embeddings), and CMonge (a conditional optimal-transport model).^6–9,12^ For the perturbation screens the baseline set was chosen to span the model families that dominate that literature, from knowledge-graph and foundation-model predictors to compositional autoencoders and neural optimal transport. Two training-free references were also scored: trainMean (the mean over training perturbed cells) and, for genetic and drug combinations, an Additive prediction of the control mean plus the two single deltas.

### Evaluation

Every held-out perturbation, combination or timepoint was scored against the corresponding observed target population in gene-expression space. Each model retained the native representation, encoder and prediction architecture of its reference implementation, and only the final outputs were harmonized for evaluation. For every task, AnnFlux predictions in the 32-dimensional DRVI latent space were mapped to log-normalized expression using a closed-form ridge readout with λ = 10 fitted on the training split. This readout was used in place of the DRVI NB decoder. For the time-course task, the baseline implementations returned predictions in different transformed representations. Predictions from PRESCIENT, scDiffEq and CellFlow were mapped back by inverse principal-component transformation, whereas ARTEMIS predictions were mapped back by inverse z-score transformation. All predictions were then represented in a common log-normalized expression space, defined as the log1p of counts normalized to the median library size, before scoring. The observed held-out timepoint and 0 h cells served as the target and control populations, respectively. In the single-perturbation and combinatorial tasks, GEARS returned a condition-level mean expression vector and scGPT returned cell-level gene-expression predictions through their native output heads. CPA and chemCPA decoded their counterfactual latent states through their learned expression decoders, whereas CMonge decoded transported latent states through its autoencoder decoder. These outputs were evaluated directly in the common log-normalized expression space without an additional post hoc readout.

### Metrics

All reported metrics compared the predicted population with the matched real target in the shared log-normalized space. We write *μ*_pred_, *μ*_obs_ and *μ*_ctrl_ for the cell-mean expression vectors of the predicted, observed and control populations and Δ_•_ = *μ*_•_ − *μ*_ctrl_ for the corresponding pseudobulk change vectors. Differentially expressed genes (DEGs) were selected per condition by a two-sided Wilcoxon rank-sum test of the observed against the control cells with Benjamini-Hochberg control of the false discovery rate; among genes with *p*_adj_ < 0.05 the *k* = 50 with the largest absolute log2 fold-change were retained (equation (9)),

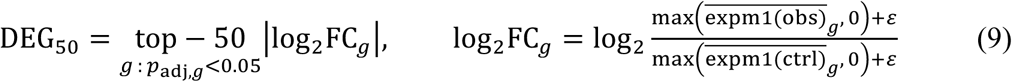

with fold-changes formed on the linear scale by inverting the log transform (expm1) before averaging. When fewer than 50 genes met the significance threshold, all significant genes were retained. A condition contributed to a top-DEG metric only when at least ten genes were significant. On the top-50 DEGs we report the shared-control-debiased Pearson correlation of the change vectors (PCC-delta top-50 DEGs), in which the control pool is split into disjoint halves *A* and *B* over 20 random repeats so that the observed and predicted Δ reference different control cells (equation (10)),

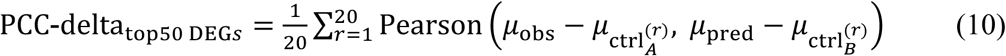

and the fraction of observed top-50 DEGs whose predicted change matched the observed sign, referred to as Common top-50 DEGs (equation (11)),

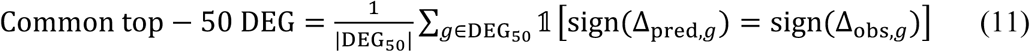

Predicted and observed populations were subsampled to a common size *m* = min(|pred|, |obs|,500) and compared by the unbiased squared maximum mean discrepancy, reported as MMD throughout, with a single median-heuristic radial-basis kernel (equation (12))^39^,

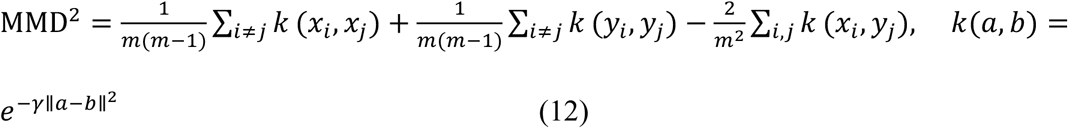

and by the Sliced-Wasserstein distance over *L* = 50 random one-dimensional projections (equation (13))^40^,

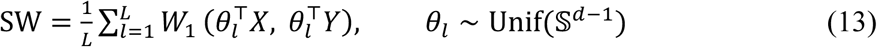

Higher PCC-delta top-50 DEGs and Common top-50 DEGs, and lower MMD and Sliced-Wasserstein distance, indicate better predictions.

### Head-to-head win rates

For each screen, metric and baseline we computed a paired per-perturbation win rate over the perturbations scored by both AnnFlux and the baseline. Scores were first averaged over the three training seeds, leaving one paired observation per perturbation. The win rate is the fraction of paired perturbations on which AnnFlux is strictly better in the metric’s own direction, that is, higher for PCC-delta top-50 DEGs and Common top-50 DEGs and lower for MMD and Sliced-Wasserstein distance. Exact ties were counted as non-wins and kept in the denominator. The two differential-expression metrics were restricted to perturbations with at least ten significant differentially expressed genes, whereas the two distributional metrics used all perturbations. Win rates were computed separately for each learned baseline, and the training-mean null was excluded. Each pairing was additionally tested by a two-sided Wilcoxon signed-rank test with zero differences discarded, corrected by the Benjamini-Hochberg procedure across all dataset × metric × baseline comparisons within each task.

### Distribution-overlay embeddings

For each held-out perturbation, a fixed reference embedding was constructed from the observed control and perturbed cells in highly variable gene-expression space using a 50-component principal-component analysis followed by UMAP (25 neighbors, minimum distance 0.4, seed 0; umap-learn 0.5.12). Each predicted population was subsampled to match the observed target cell count and projected into this reference embedding using the fitted PCA and UMAP transformations, neither of which was refitted on predicted cells. GEARS was omitted because it does not produce cell-resolved predictions. Exemplar perturbations were selected using a prespecified maximin ranking procedure. Candidates were required to have at least 50 cells and a control-versus-perturbed SNRd in the upper half of the within-screen distribution, where SNRd was defined as the mean of the twenty largest absolute Cohen’s d values across genes. Among candidates predicted by all models and having at least ten significant DEGs, PCC-delta top-50 DEGs and Common top-50 DEGs were converted to within-model percentile ranks. The selection score was the minimum percentile rank across both metrics and all models, such that a perturbation ranked highly only when it was consistently ranked highly by every model on both metrics.

The time-course overlays followed the same principle, but each reference embedding spanned the complete measured trajectory. Expression was recomputed from raw counts as median-library-size-normalized log1p values over 2,000 highly variable genes. A 50-component principal-component analysis followed by UMAP (25 neighbors, minimum distance 0.4, seed 0) was fitted to all observed cells at 0 h, 8 h, 1 d, 3 d and 7 d. Predictions from every model were projected using the fitted PCA and UMAP transformations, neither of which was refitted on predicted cells.

### Object-wise drift decomposition

To characterize inducer-specific drift, we evaluated the trained model with one object slot active at a time (**Fig. 2a and b**). The drift assigned to each inducer was defined as the complete conditional drift evaluated with that inducer alone. This analysis used the EMT model on A549 under the cell holdout, taking the source pool from the held-out test cells.^3^ For each inducer the per-object drift was averaged over 400 of these cells at 0 h on a normalized time grid t ∈ {0.1, 0.3, 0.5, 0.7, 0.9}, and its magnitude taken as the L2 norm of the latent drift. As a leakage control, the 70/10/20 split (seed 42) was reproduced and inducer drifts were computed independently from the train, validation and test 0 h cells, comparing train-versus-test cosine and magnitude. To represent these drifts in gene space, each inducer-isolated drift was integrated deterministically (noise-free Euler, 30 steps) from z₀ to z_end_ and mapped to log-normalized expression by the ridge decoder, giving Δ = decode(z_end_) - decode(z₀). This endpoint is not the conditional mean of the stochastic dynamics. On the same source cells the two agreed in decoded direction with cosine 0.92 to 1.00, and their norms differed by at most 12%. Drift direction was assessed in this gene space, comparing the per-inducer Δ vectors by pairwise cosine and projecting the three jointly by two-dimensional principal component analysis.

Inducer specificity was assessed by a token-swap counterfactual on a single fixed pool of cells (**Fig. 2c**). From the held-out test split we took up to 300 A549 cells at 0 h and, holding this source pool and its latent coordinates fixed, integrated the drift field once under each of the three inducer tokens in turn, so that any difference between conditions is attributable to the object token alone. Integration was deterministic (noise-free Euler, 30 steps, midpoint time sampling) and each endpoint was mapped to gene space by a ridge readout (λ = 10); the decoded drift is the difference between the mean decoded expression at the endpoint and at the 0 h source, giving one signed value per highly variable gene.

### Program-level analysis of decoded drifts

Hallmark gene-set enrichment of this drift direction used gseapy^41^ prerank against MSigDB Hallmark 2020^42^ (10,000 permutations), reporting the normalized enrichment score and FDR q-value of the EMT term.

For the pathway-specificity matrix (**Fig. 2c**), we performed a token-swap counterfactual analysis on a fixed held-out 0 h source pool. The decoded drift vector was used directly as the ranking metric for a preranked gene-set enrichment analysis (gseapy prerank, 1,000 permutations, minimum set size 5, maximum 1,000, seed 42) against the MSigDB Hallmark 2020 collection.^42^ Four terms are reported: TGF-beta Signaling, TNF-alpha Signaling via NF-kB, KRAS Signaling Up (used as a RAS/MAPK proxy because no EGF-specific Hallmark set exists) and Epithelial Mesenchymal Transition. Each cell reports the mean normalized enrichment score across three training seeds; asterisks indicate an FDR q value below 0.05 in all three seeds. The matrix displays four of the 48 Hallmark terms represented by at least five genes.

EMT master-regulon activity was scored with AUCell over the CollecTRI network for five core regulators (*SNAI2*, *ZEB1*, *ZEB2*, *SNAI1* and *TWIST1*).^43,44^ For each inducer, the change in activity from 0 h to the drift-driven endpoint was reported as the mean over three seeds. TF alignment scored the mean Δ over the up- versus down-regulated members of the conserved Cook EMT regulon (up: *JUN*, *JUNB*, *FOS*, *RELB*, *ATF4*, *SOX4*, *KLF6*; down: *ELF3*).

### Program axes and drug-program decomposition

Lineage and drug-response program axes are individual DRVI latent dimensions selected by a marker-panel correlation screen rather than by decoder loading rank. For each curated marker panel a per-cell program score was computed as the mean z-scored highly variable gene expression over the panel genes, the Pearson correlation of every latent coordinate with every program score was evaluated across all cells, and the dimension of largest absolute correlation was assigned to that program. On the Norman K562 model this gave latent dimension 23 for the erythroid program^22,23^ (Pearson r = +0.84) and dimension 30 for the myeloid program (Pearson r = -0.76). Each axis was oriented so that increasing values correspond to increasing program activity, by taking the sign of the mean ridge-decoder loading over that dimension’s marker panel; the axis magnitude is the latent coordinate itself. A perturbation was projected onto these axes by integrating the drift field deterministically from control cells (noise-free Euler, 20 steps, 300 control cells) and reading the latent coordinate along integration time t, averaged over three seeds, with each TF decoded only by the folds in which it was held out. For the lineage-commitment trajectories the reported quantity is the oriented erythroid-minus-myeloid contrast with the control value at t = 0 subtracted, so that every perturbation starts at zero.

### Pathway recovery and set overlap

To test whether a predicted response reproduces the observed transcriptional program, we compared the pathways over-represented among each population’s top response genes. The query gene set was the 150 genes with the largest absolute mean shift relative to the shared control mean, taken from the predicted or the observed population, and the background was the dataset’s highly variable gene panel. Over-representation against the Reactome 2022 pathway collection from the Enrichr gene-set libraries was tested per pathway by a one-sided Fisher exact test and corrected across pathways by the Benjamini-Hochberg procedure, calling a pathway enriched at a false-discovery rate below 0.05.^45,46^ Agreement between the predicted and observed enriched-pathway sets was the Jaccard index of their significant term sets, computed for perturbations with at least five significant observed pathways; the significance of a given overlap was assessed by a hypergeometric survival probability using the tested Reactome terms as the universe, with Benjamini-Hochberg correction across each screen’s held-out perturbations. For the whole-screen comparison, each model’s per-perturbation Jaccard was tested against AnnFlux by a one-sided paired Wilcoxon signed-rank test with Benjamini-Hochberg correction.

### Prior-knowledge ablation and drift-space organization

The contribution of the prior-knowledge network was measured by a paired ablation in which the prior encoder was replaced with a learned object-embedding encoder with the same interface, removing the STRING and LINCS-L1000 channels together, and the model was retrained under an otherwise identical protocol. For each dataset, perturbation regime and metric, the score of every held-out perturbation was averaged over the three training seeds within its evaluation fold. The paired effect size was calculated as Cohen’s dz after orienting all metric differences so that positive values indicated better performance with the prior. Significance was assessed using a paired Wilcoxon signed-rank test with Benjamini-Hochberg correction. The perturbation is the unit of analysis: seed-fold repeats re-measure the same perturbations and are not independent replicates of the prior’s effect.

Drift-space organization was assessed using 104 Norman single-gene perturbations. For each perturbation, the 32-dimensional DRVI drift response was defined as the endpoint shift from control after integrating the learned SDE from control cells. Responses were averaged across the five fold-specific models. The resulting matrix was standardized feature-wise and centered, and its first two singular components, representing dominant variation shared across perturbations, were removed before cosine similarities were calculated. Recovery of known biological groupings was quantified by the AUPRC for ranking within-group pairs above between-group pairs. The analysis included 32 genes from 10 HGNC gene families and nine genes from three K562 lineage programs: erythroid, myeloid and megakaryocytic. We compared the no-PKN model, the STRING-prior model and the STRING Node2Vec embedding. Significance was assessed using a label-shuffling null with 5,000 permutations. Chance-level AUPRC, given by the positive-pair fraction, was 0.085 for gene families and 0.278 for lineage programs.

### Additive baseline and genetic-interaction classes

For a combination of perturbations A and B, the additive prediction was the sum of the two observed single-perturbation mean changes, Δ_add_ = Δ_A_ + Δ_B_, where Δ_X_ is the difference between the mean log-normalized expression of cells receiving perturbation X and control cells over the ∼2,000 highly variable genes. The vector-field panels read one combination model at a single integration time. For each single perturbation the drift at t = 0.5 was taken relative to the drift under the control token, F(A) - F0, and projected onto the erythroid and myeloid axes. Projected drifts were averaged as unit vectors within a 10 × 12 grid and drawn at fixed length. Bins holding fewer than five cells were dropped. The predicted combination cloud was integrated from held-in control cells by noise-free Euler over 30 steps.

Each combination’s genetic-interaction class was taken directly from the GEARS genetic-interaction annotation (the GIs dictionary in gears/inference.py), which labels combinations with the five non-additive interaction types defined by Norman et al. together with an additive class.^8,22^ This gold-standard set comprised 88 two-gene Norman combinations: 30 synergistic, 16 additive, 13 neomorphic, 12 suppressor, 9 epistatic and 8 redundant.

### Cross-modality evaluation against CD11b surface protein

Predictions were evaluated against protein measurements from the paired CITE-seq combinatorial CRISPR screen of Wessels et al.^24^. CD11b antibody-derived tag (ADT) counts were centered-log-ratio normalized^47^ by applying log1p followed by per-cell centering across the antibody panel. Each perturbation was summarized as the difference between its mean CD11b value and the control mean. Only perturbations with at least 10 cells were included, yielding 158, 129 and 63 combinations in the 2 seen, 1 seen and 0 seen strata.

A monocyte and macrophage differentiation program was taken from the MSigDB C5 GO biological process collection (v2024.1.Hs)^42,48^: the union of GOBP_MONOCYTE_DIFFERENTIATION (GO:0030224) and GOBP_MACROPHAGE_DIFFERENTIATION (GO:0030225) (up) minus GOBP_HEMATOPOIETIC_PROGENITOR_CELL_DIFFERENTIATION (GO:0002244) (down). Genes shared between the two components, all perturbed target genes and genes outside the highly variable set were removed, leaving 17 up-genes and 22 down-genes. In particular the set does not contain ITGAM, the gene encoding CD11b, so the predicted RNA program and the measured ADT readout share no gene. The program score was the mean expression change over the up-genes minus the mean over the down-genes, computed from each combination’s mean shift relative to control.

Within each stratum, cross-modality agreement was the Spearman correlation across combinations between the predicted program score and the observed CD11b shift. Correlations were computed only when at least five paired combinations remained and the predicted program score was not constant across the stratum. Correlations were reported as the mean over three seeds (42, 1024 and 2026), with a 95% confidence interval from a bootstrap over combinations. For the 2-seen stratum, the ceiling and additive references were computed using the 142 combinations retained for that seed; the additive reference was defined as the sum of the two observed single-gene responses.

### Spatial projection of the IFN-response signature

The Frangieh screen was performed under IFN-γ stimulation, providing a context for identifying perturbation responses associated with IFN-γ signals.^27^ We focused on STAT1 and JAK2, as two canonical components of this pathway at complementary signal levels: STAT1, the TF shared by type-I and type-II IFN signaling^27^, and JAK2, the kinase required for IFN-γ receptor signaling.^28^ We derived an interferon-response signature from AnnFlux predictions on the Frangieh melanoma screen under IFN-γ stimulation (three seeds, five folds). STAT1 and JAK2 were treated as held-out perturbations: each gene’s contribution was read from the cross-validation fold in which it was the evaluation perturbation, so the signature reflects predicted rather than memorized knockout effects. For each gene we integrated the object-conditioned drift noise-free over 30 steps from 300 held-out control cells carrying that gene’s knockout token under the IFN-γ condition, decoded the endpoint through the ridge readout and took the gene-space shift relative to the source, averaged over the three seeds. The two genes’ shifts were averaged and the signature was defined as the negated shift, so that a positive weight marks a gene suppressed by knockout, that is, an IFN-γ-induced gene; the 100 genes with the largest absolute weight were retained and used for all downstream spatial scoring.

We projected the signature onto the Cho pan-cancer spatial atlas of tertiary lymphoid structures (TLSs)^25^ (35 Visium sections across eight cancers; sections with fewer than 30 spots excluded). Each spot received a signed signature score, the weight-normalized weighted mean of the per-gene z-scores of log1p expression: with signature weights w and per-gene mean μ and standard deviation σ computed across spots, the score equaled the sum of w(x - μ)/σ over the signature genes divided by the sum of |w|, where x is the spot’s log1p expression. The 107 atlas programs, comprising curated meta-programs and MSigDB Hallmark gene sets, were represented by their atlas-provided AUCell scores.^42^ Distance from intratumoral TLSs was represented by the atlas-provided normalized per-spot distance. For each section we computed the Pearson correlation between the per-spot signature score and this distance. The sections shown in **Fig. 5a and Fig. 5b**, one kidney and one liver, were the two sections with the strongest negative correlations among sections from distinct cancer types. Following Cho et al.,^25^ we characterized spatial gradients by fitting each program score as a natural cubic spline of normalized TLS distance with three degrees of freedom. The fitted curve was evaluated on a 100-point grid and z-scored, and gradient strength was defined as the proximal-minus-distal endpoint difference. A program was classified as high-to-low when the fitted curve was non-negative at the proximal endpoint and non-positive at the distal endpoint, low-to-high under the reverse pattern and mixed otherwise. Within each cancer, gradient significance was assessed by comparing the spline with an intercept-only model using an F-test, followed by Benjamini-Hochberg correction across programs. The distance profiles compare the AnnFlux signature with the Hallmark IFN-α and IFN-γ programs in cancers where the Hallmark IFN-γ program showed a significant high-to-low gradient (**Fig. 5c**). At the section level, a positive proximal-minus-distal difference was classified as TLS-proximal and a non-positive difference as TLS-distal.

### Statistics

Unless stated otherwise, n denotes the number of perturbations or combinations scored, and learned models were run over three training seeds (42, 1024, 2026). Metrics were computed per perturbation and seed, pooled within a seed across folds, and summarized as the mean ± standard deviation across seeds; deterministic baselines report a single value, and top-DEG metrics were averaged only over perturbations with at least ten significant DEGs. Differential expression used a two-sided Wilcoxon rank-sum test (Mann-Whitney U) with Benjamini-Hochberg correction (q < 0.05). For the single-perturbation task, AnnFlux was compared with each baseline by the same paired two-sided Wilcoxon signed-rank test, run separately for each metric and screen. No statistical method was used to predetermine sample size, and model evaluation was not performed under blinding.

## Data availability

The datasets used in this study are publicly available. The Cook time course is available at GEO accession GSE147405; the Replogle, Norman, Frangieh, sci-Plex3 and Wessels screens were obtained as processed single-cell objects from the scPerturb collection^49^ (Zenodo 13350497); and ComboSciPlex was obtained from the CPA distribution (Lotfollahi et al. 2023)^6^ (GEO accession GSE206741).

## Code availability

All source code, preprocessing scripts, trained model weights, benchmark outputs and figure-generation scripts will be made publicly available upon submission at https://github.com/joonan-lab/AnnFlux. Models were implemented in PyTorch, with scanpy^37^ for preprocessing, DRVI^38^ for the latent autoencoder, gseapy^41^ for gene-set enrichment and RDKit^50^ for molecular fingerprints.

## Acknowledgements

We are grateful to Hwanseok Sim for his assistance with the model architecture.

## Author contributions

Conceptualization: JYA

Methodology: HSC, GEB, HYP, SBL, JYA

Software: HSC, GEB, HYP, JYA

Formal analysis: HSC, GEB, HYP, JSP, JYA

Data curation: HSC, GEB, HYP, JYA

Writing - original draft: HSC, GEB, HYP, JYA

Writing - review & editing: HSC, GEB, HYP, JSP, SBL, JYA

Visualization: HSC, GEB, HYP, JSP, JYA

Funding acquisition: JYA Supervision: JYA

Project administration: JYA

